# Quicklime-based eradication attempt of *Xenopus laevis* as a model for controlling pondscape invasions

**DOI:** 10.64898/2026.02.14.705882

**Authors:** Teun Everts, Loïc van Doorn, Tim Adriaens, Jeroen Speybroeck, Nicolas Pardon, Marco Morbidelli, Sabrina Neyrinck, Johan Auwerx, Lia Baeteman, Maud Segal, Rein Brys

**Affiliations:** Research Institute for Nature and Forest, Brussels, Belgium; KU Leuven, Heverlee, Belgium; Agency for Nature and Forest, Brussels, Belgium; University of Florence, Florence, Italy; Ghent University, Gent, Belgium

**Author notes:** Corresponding author: Teun Everts.

**Keywords:** African clawed frog, biological invasions, calcium oxide (CaO), species removal, early-detection-rapid-response, management costs, non-native invasive species

## Abstract

Aquatic non-native invasive species are notoriously difficult to eradicate, particularly in pondscapes where populations can spread rapidly, persist in unmanaged refugia, and recolonise treated sites. In such contexts, high-intensity management interventions may be justified, balancing short-term collateral impacts against the prevention of permanent establishment and long-term damages. Chemical eradication methods, such as rotenone or herbicide application, can be effective but raise ethical and environmental concerns. Here, we evaluate quicklime (calcium oxide, CaO) application as a more sustainable alternative control tool for pondscape invaders compared to other chemical methods, using the African clawed frog (*Xenopus laevis*) invasion in Belgium as a case study. When applied to water, quicklime hydrates exothermically to calcium hydroxide (Ca(OH)), which releases OH ions upon dissolution, temporarily and rapidly increasing pH to lethal levels. In winter 2023, three ponds with breeding populations of *X. laevis* of low ecological value were drained, fenced, and treated with quicklime. Treatment effectiveness was assessed through pH measurements, visual surveys, and environmental DNA (eDNA) quantification. Immediately after treatment, large numbers of deceased post-metamorphic individuals were recovered, indicating treatment-induced mortality. Eight weeks post-treatment, eDNA concentrations were markedly lower in two of the three ponds (reductions of 100% and 80%) compared to those during the same period one year later. Although eDNA concentrations increased again during the following summer suggesting partial population recovery through survival and/or recolonisation, they remained lower than pre-treatment conditions. Water pH returned to near baseline levels within one month. We provide the first field-based preliminary evidence that quicklime can induce large-scale mortality in *X. laevis* populations in small to medium-sized ponds. We discuss practical considerations, limitations, and broader applicability, proposing quicklime as a high-intensity option within integrated management strategies for pondscape invaders.

## Introduction

Countries across the globe are experiencing adverse effects of non-native invasive species (Gómez-Suárez al., 2025), yet the expectation is that more will occur in the coming decades (Seebens et al., 2017; 2025). This trend is of particular concern because the introduction, establishment, spread (Haubrock et al., 2025a), and impact (Haubrock et al., 2025b) of non-native species, collectively referred to as biological invasions, are increasingly recognised as major threats to biodiversity and global sustainability (IPBES, 2023). Aquatic ecosystems are particularly susceptible to biological invasions (Reid et al., 2018; Cahill et al., 2026) due to characteristics such as low species detectability, particularly during early invasion stages (Harvey et al., 2009; Measey et al., 2012; Simberloff, 2021), and strong trophic links that can be disturbed by the loss or the introduction of species (Gallardo et al., 2016).

Once established, aquatic non-native invasive species are often difficult to eradicate (Pluess et al., 2012; Simberloff, 2021; Cahill et al., 2026). The high level of connectivity of aquatic systems not only hampers removal up to the very last individual, but also facilitates rapid recolonisation from nearby source populations, undermining eradication efforts (Britton et al., 2008; Cahill et al., 2026). In pondscapes, defined as a network of ponds and their surrounding terrestrial habitat matrix (Boothby, 1997), non-native invasive species can spread rapidly, persist in unmanaged refugia, and recolonise managed sites, facilitated by *via* canals, drainage ditches, flood events, or overland dispersal (Simberloff, 2021). These dynamics pose particular difficulties for managing non-native invasive amphibians, whose complex life cycles spanning aquatic and terrestrial habitats allow them to evade site-based interventions and recolonise treated systems (Vimercati et al., 2017; Everts et al., 2022). Consequently, rapid response to incipient aquatic invasions, ideally before measurable impact occurs, using highly effective interventions, are critical to successful management of pondscape invaders (Reaser et al., 2020; Simberloff, 2021; Spear et al., 2021; Everts et al., 2025a).

If not limited by legal restrictions, management is in such contexts often framed around the notion that “*the ends justify the means*,” whereby short-term collateral impacts are weighed against prevention of permanent establishment and the associated ecological and economic damages (Pluess et al., 2012; Green & Grosholz, 2020; Diagne et al., 2021; Simberloff, 2021; Haubrock et al., 2025c). Chemical eradication represents one such approach, which commonly relies on introducing chemical compounds to the water, such as pesticides, including rotenone or antimycin A for fish (Rytwinski et al., 2019), herbicides for macrophytes (Kujawa et al., 2017), caffeine and chloroxylenol for amphibians (Witmer et al., 2015), and Virkon S (DuPont Inc.) at for fungi (Bosch et al., 2015). Although these methods are often effective and cost-efficient (Britton et al., 2008; Rytwinski et al., 2019), they raise concerns regarding non-target impacts, incomplete removal requiring repeated treatments, the evolution of resistance, environmental persistence of chemical compounds, and the risk of spill-over into uninvaded systems (Kujawa et al., 2017; Dunlop et al., 2018; Beaulieu et al., 2021; Kjærstad et al., 2021).

An often overlooked but potentially effective eradication measure in pondscape invasion management is the application of quicklime (CaO) (Clearwater et al., 2008). Quicklime and other lime-based treatments such as slaked lime, and dolomite have a long history of use as a disinfectant in aquaculture and waterbody sanitation. Upon contact with water, CaO hydrates exothermically to calcium hydroxide (Ca(OH)h). Dissolution of Ca(OH)h releases Ca2⁺ and OH⁻ ions (Ca(OH)h ri Ca2⁺ + 2OH⁻), rapidly increasing alkalinity and driving pH to levels that can be lethal to most aquatic life (Boyd, 2017). Over time, uptake of atmospheric CO and subsequent carbonate equilibria consume OH ions, promoting bicarbonate and carbonate formation and the precipitation of calcium carbonate (CaCO), thereby allowing pH to gradually return toward pre-treatment conditions.

In aquaculture, quicklime and other lime-based treatments are widely applied to regulate pond productivity, to improve nutrient availability and buffering capacity, and to remove unwanted organisms and disease vectors between production cycles (Wurts & Masser, 2013; Boyd, 2017; Lazur et al., 2002). This broad-spectrum mode of action has also been explored beyond aquaculture, including sea urchin culling to promote kelp recovery in Norwegian fjords (Christie et al., 2024) and sea lice control in salmon farming (Brooks et al., 2020). In the context of invasion management, the principal advantage of quicklime lies in its capacity to achieve rapid, whole-system population removal, which can be particularly effective in small and relatively closed aquatic systems. This may reduce the likelihood of compensatory, density-dependent responses, such as increased growth, survival, or recruitment following partial population suppression (Govindarajulu et al., 2005), as well as evolutionary responses that can undermine repeated or incomplete control efforts (Dunlop et al., 2018; Siddiqui et al., 2023; Zarri et al., 2025). Quicklime is highly effective in killing aquatic organisms. The optimum pH range for most aquatic organisms is 6.5–8.5, and the acid and alkaline death points are around pH 4 and pH 11, respectively (Boyd, 2020). Most fish, invertebrates, and amphibians cannot survive prolonged exposure to water with pH levels above 12, resulting in rapid mortality. Also, in systems with medium to high pH, liming precipitates calcium phosphate and calcium carbonate which removes phosphorus from the water column. This promotes sedimentation and is sometimes applied as eutrophication control as it binds phosphates and nitrogen, reducing nutrient availability and thereby avoiding rapid proliferation of algae and decreases in dissolved oxygen linked with nutrient enrichment (Murphy and Prepas 1990; Yang et al. 2025). Moreover, quicklime acts through acute physicochemical alteration and its effects on water acidity are typically short-lived, with breakdown products consisting of naturally occurring minerals (Piper et al., 1982; Boyd, 2017). However, it comes at the cost that it also (lethally) affects non-target species (Abdel-Fattah, 2011). Explicit consideration of non-target impacts and post-treatment recovery trajectories is therefore essential when evaluating its suitability as a management tool.

Here, we use the invasion of the African clawed frog (*Xenopus laevis*) in Belgium to assess whether quicklime application could provide a sustainable and effective alternative to other chemical approaches for eradicating pondscape invaders. We describe the application and evaluation of a quicklime treatment to three *X. laevis* breeding ponds within the framework of an early-detection-rapid-response programme aimed at preventing permanent establishment of this species and its associated pathogens in Belgium. More specifically, we provide preliminary evidence for the effectiveness of quicklime in eradicating *X. laevis* populations in both the short term (weeks after treatment), and the medium-term (months after treatment), and evaluate the advantages and limitations of the technique, including its potential applicability to other non-native pondscape invaders.

## Methods

### Study species

The African clawed frog (*Xenopus laevis,* Fig. 1) is a primarily aquatic amphibian species, native to Sub-Saharan Africa. This opportunistic anuran species tolerates a broad spectrum of environmental conditions, and is thus able to colonise a wide variety of water bodies (Measey et al., 2012). Breeding mainly occurs in both temporary and permanent ponds, although it can also occur in slowly flowing lotic systems (Moreira et al., 2017). The species survives extreme conditions through entering a dormant state buried into mud or pond sediment (Measey, 2016). Migration can occur overland (Measey, 2016), but is facilitated by ditches, streams, and rivers (Vimercati et al., 2024; Everts et al., 2025a).

**Figure 1.**
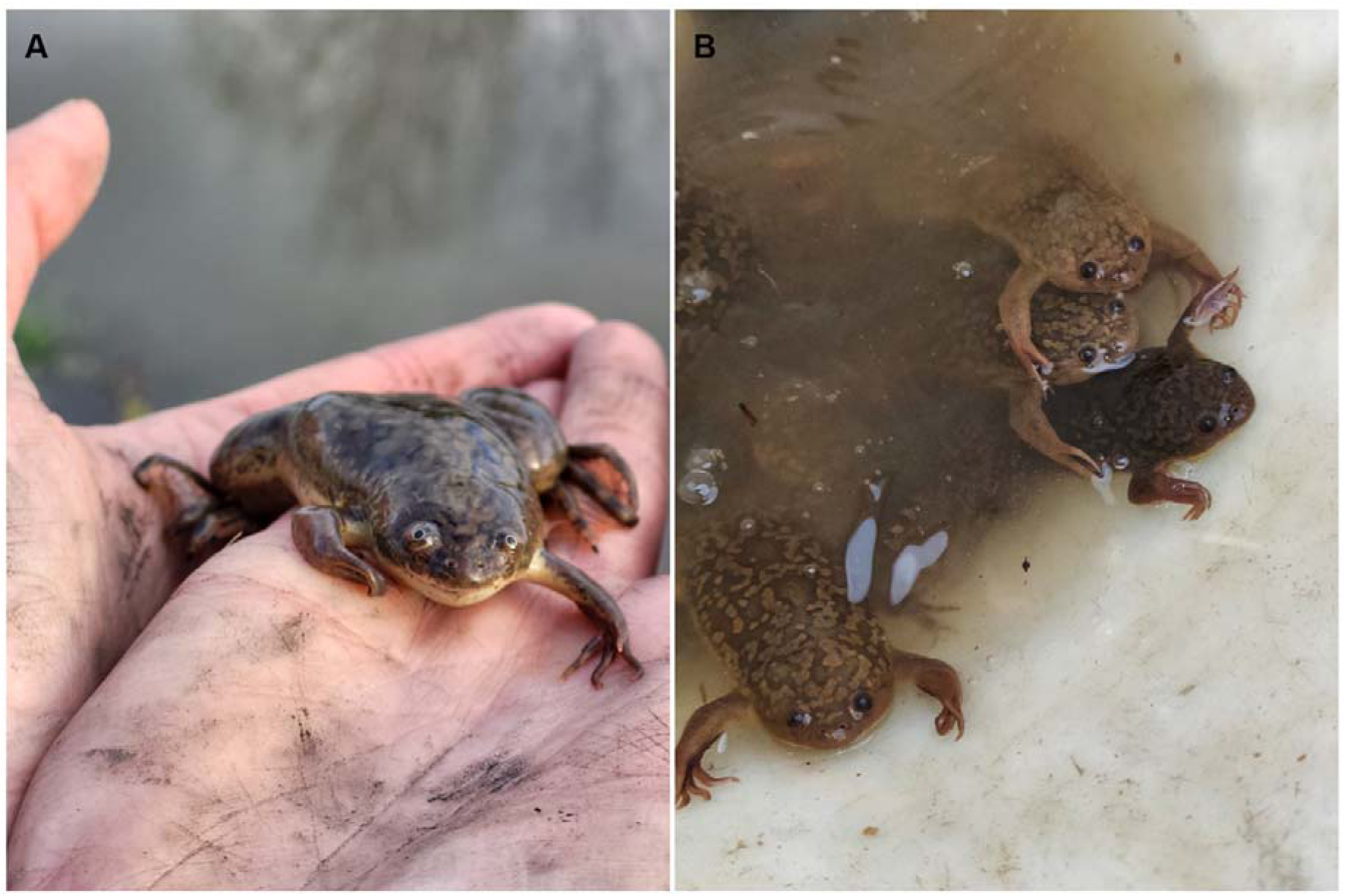
Target species used to evaluate the effectiveness of quicklime for eradicating non-native invasive species in pondscapes: the African clawed frog (*Xenopus laevis*). (**a**) Post-metamorphic individual; (**b**) multiple individuals captured at the study sites prior to quicklime treatment.

Through the pet trade and its use as laboratory animals, *X. laevis* was introduced outside its native range. Established non-native populations now occur in North America, South America, Asia, and Europe (Measey et al., 2012; Sittert & Measey, 2016). Although *X. laevis* introductions started in the 20^th^ century, new populations continue to be discovered worldwide (van Doorn et al., 2022; Pauwels et al., 2023; Emmenegger et al., 2025; Everts et al., 2025a). This is worrying, as *X. laevis* can reach high densities (> 13 frogs/m² Mora, 2019) and exert strong ecological impacts on recipient environments. *Xenopus laevis* is characterised by ontogenetic dietary shifts. Whereas tadpoles are nektonic, phytophagous filter-feeders (Schoonbee et al., 1992), metamorphosis leads to bottom-dwelling predators mainly feeding on zooplankton, zoobenthos, amphibians, and invertebrates (Amaral & Rebelo, 2012; Courant et al., 2018). Their toothed jaws and fast hindlimb movements enable *X. laevis* to catch and shred large-sized prey (Avila & Frye, 1978; Measey, 1998a). As such, *X. laevis* can reduce food availability in and around aquatic systems, trigger trophic cascades, and initiate fierce resource competition with native species (Courant et al., 2018; Measey, 1998b). Their bottom-dwelling behaviour increases water turbidity and nutrient release, thereby deteriorating water quality (Lobos & Measey, 2002). Moreover, similar to other non-native amphibians (Everts et al., 2025b), *X. laevis* can be a carrier of the chytrid fungus (Schmeller et al., 2011), one of the biggest threats to amphibians worldwide (Scheele et al., 2017), a potential vector of *Ranavirus* (Robert et al., 2007), and carrying a diverse range of other parasites and pathogens including tapeworms, nematodes, and ciliates (Schoeman et al. 2019, 2020). Clearly, *X. laevis* is a highly invasive non-native species and was therefore added to the list of non-native species of European Union (EU) concern under EU Regulation 1143/2014. As such, EU member states are obliged to undertake management measures, including early-detection-rapid response interventions, to combat this species.

### Study system

The applicability of quicklime for the eradication of *X. laevis* in Belgium was evaluated in three ponds where the species was first detected and confirmed to be breeding (Fig. 2). Here, rapid eradication of *X. laevis* was primarily considered to prevent its potential permanent establishment, yet precautionary measures were also intended to limit the potential establishment of associated pathogens (Robert et al., 2007; Britton et al., 2008; Hossack et al., 2023). Dipnetting and fyke netting in the summer of 2022 confirmed that *X. laevis* was present in high densities and in all life stages in each of these three ponds, indicating breeding activity (Pauwels et al., 2023). All three ponds are located within the Douvebeek river valley, within 50 metres from the river (Fig. 2). The Douvebeek is a small, turbid stream approximately 21 kilometres in length flowing along the French-Belgian border, characterised by a highly variable precipitation-driven discharge regime, ultimately flowing into the river Lys. The surrounding landscape is dominated by intensive agriculture interspersed with small towns. Soils are predominantly sandy loam to loam, with locally occurring alluvial loam–clay soils. Despite having ecological value in its upper reaches, the Douvebeek is overall in poor condition and has significant water quality issues due to discharge, erosion, and agricultural runoff.

**Figure 2.**
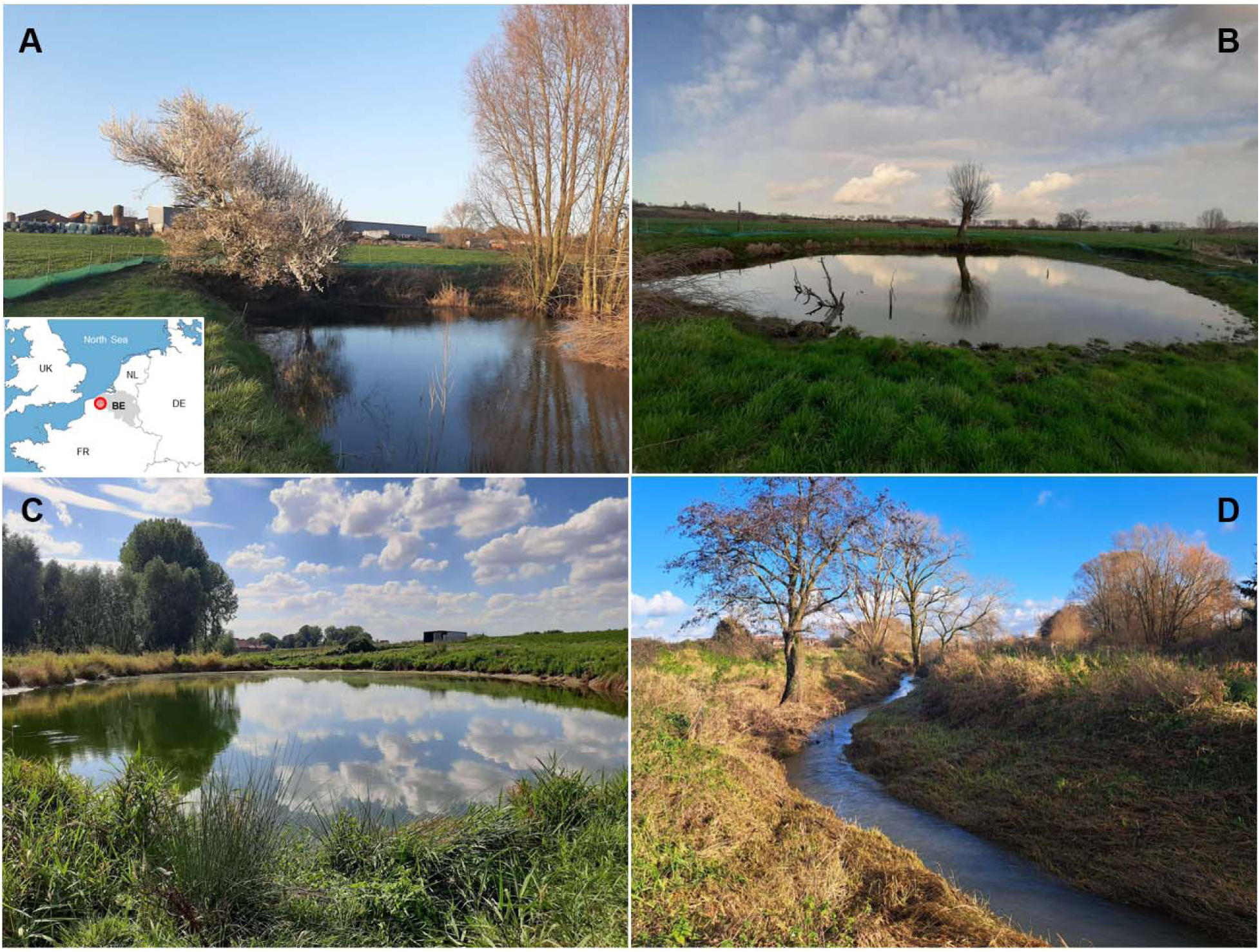
The three ponds that were subject to a quicklime treatment during the winter of 2022-2023 to eliminate locally breeding *Xenopus laevi*s populations: (**a**) Pond 1 (50° 45’ 19.69” N, 2° 53’ 9.45” E) together with a map showing the geographic location of the study area, (**b**) Pond 2 (50° 45’ 18.84” N, 2° 53’ 10.60” E), and (**c**) Pond 3 (50° 45’ 24.00” N, 2° 55’ 6.19” E). All ponds are situated within 50 metres of the Douvebeek river (**d**), illustrated here during a low-flow period; river volume can increase by up to 200% following intense precipitation events.

Pond 1 (surface area: 600 m²) functions as a water-retention pond for irrigation, exhibiting pronounced fluctuations in hydroperiod. Its large depth, exceeding 3 metres at its deepest point, reduces the likelihood of complete desiccation. The pond is surrounded by tree and shrubs that cast substantial shade, and aquatic macrophytes are largely absent. Pond 2, located between the Douvebeek and Pond 1 (Fig. 2), is a small (480 m²) cattle-drinking pond with a maximum depth of 1.5 metres. While it is fully exposed to sunlight and periodically supports extensive submerged macrophyte growth, it is highly disturbed by cattle through trampling. Pond 3, approximately 2 kilometres located from Ponds 1 and 2, is a large (1,200 m²) water-retention pond with an average depth of 2 metres, located next to agricultural fields. It is fully exposed to sunlight and is characterised by abundant riparian vegetation but limited submerged, emergent, or floating vegetation cover. The pond’s hydroperiod is strongly influenced by agricultural activity, as intensive use of water for crop irrigation reduces water availability, and during dry periods water is pumped from the Douvebeek into the pond, except when abstraction is prohibited during extreme drought conditions. All three ponds can be considered as having low ecological value due to their eutrophic, disturbed, and agricultural nature (Pauwels et al., 2023; López-de Sancha, et al., 2025).

### Treatment

Between January 10^th^ and 19^th^ 2023, all three ponds were treated with quicklime (calcium oxide; CaO) as part of an early-detection-rapid-response strategy aimed at preventing permanent establishment of *X. laevis* and mitigating associated pathogen risks (Everts et al., 2025a). Treatment was carried out during winter because *X. laevis* is expected to overwinter in ponds (Measey, 2016), and because this period falls outside the breeding period of most native amphibians, thereby aiming to minimise collateral impacts.

Prior to treatment, each pond was enclosed to prevent *X. laevis* from dispersing away from the pond in response to the drainage and quicklime treatment (Fig. 3). To this end, a 1-m-high insect-mesh barrier (mesh size: 1.0 × 1.4 mm) was installed along the perimeter of each pond and partially buried to prevent escape *via* burrowing. To further inhibit climbing or jumping individuals, the upper edge of the mesh was folded inward toward the pond. At 10-m intervals along the barrier at the side of the pond, pitfall traps (buckets buried so their upper rims were flush with the ground) were installed to intercept *X. laevis* as well as other animals moving along the fence line. To minimise the volume of water to be treated with quicklime, ponds were drained as much as possible using a wastewater pump (Clearwater et al., 2008; Witmer et al., 2015; Lazur et al., 2002) (Fig. 3). Subsequently, fyke nets were deployed in the remaining water for five consecutive days to remove as many organisms as possible prior to treatment. *Xenopus laevis* individuals were euthanised using MS-222, and native species were translocated to a neighbouring, untreated pond.

**Figure 3.**
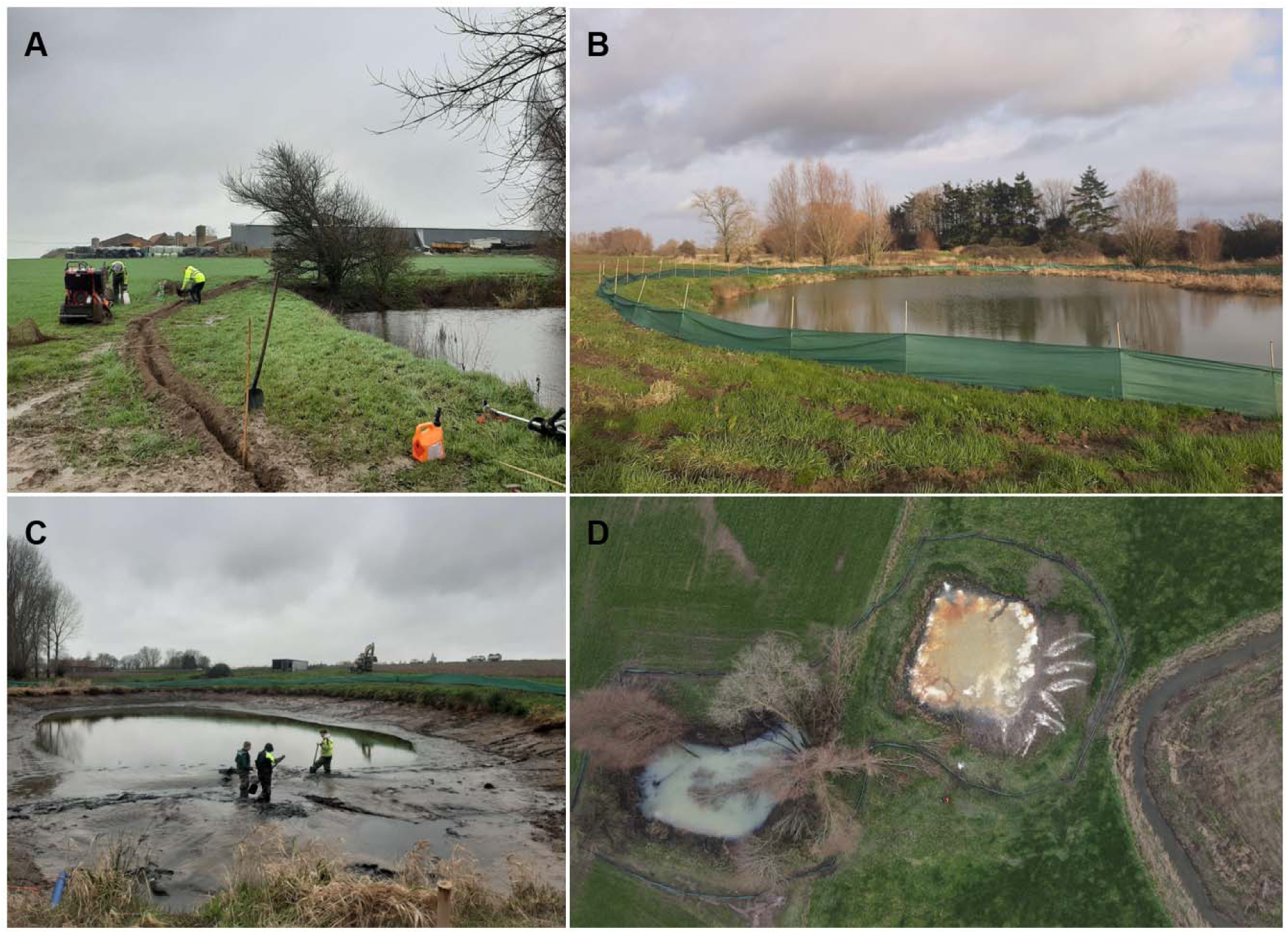
Key steps in applying quicklime to eradicate a non-native amphibian species from ponds. (**a**) Trenches, approximately 30 cm deep, were excavated using a trench cutter along the entire perimeter of the pond, and (**b**) a 1-m-high mesh barrier was installed to prevent animals from escaping the pond following treatment, including burrowing beneath the fence. (**c**) The pond (Pond 3 is shown) was drained as much as possible to reduce the water volume requiring treatment and thereby minimise the amount of quicklime needed to achieve an application rate of 0.6–3.1 kg ml³. (**d**) Appearance of Pond 1 (left) and Pond 2 (right) during quicklime treatment. The fences surrounding the ponds, as well as the Douvebeek are also shown (picture courtesy of Emilie Gelaude).

The ponds were then treated with quicklime (16^th^ of January 2023) at a minimum dosage of 0.6–3.1 kg m ³ (Dykzeul & Young, 2001; IFS 2003a, 2003b; Barnes, 2005; Wurts & Masser, 2013) (Fig. 3). A crane was used to distribute quicklime across the pond and to mix it with the remaining pond water. As effective eradication generally requires maintaining a pH above 12 for at least four consecutive days (Clearwater et al., 2008), water acidity (pH) was continuously measured and additional quicklime was applied when pH dropped below 12 during this period. To prevent recolonisation, fences were maintained for four weeks after treatment, yet subsequently removed because access to the ponds was required by landowners.

### Management evaluation

#### Quantitative eDNA-based evaluation

We evaluated the effectiveness of the quicklime treatment using environmental DNA (eDNA) barcoding on a droplet digital PCR (ddPCR) platform. In each pond, we collected three eDNA samples at three different time points: during the summer of 2022 (i.e. baseline conditions), eight weeks after treatment during the winter of 2023 (i.e. February), and during the summer of 2023 (i.e. August; short-term efficacy). To disentangle phenology-related natural seasonal fluctuations in *X. laevis* eDNA signals in summer and winter (Everts et al., 2021) from eDNA concentration drops due to quicklime treatment, we additionally conducted eDNA sampling in each treated pond in February of the year following quicklime application (2024; medium-term efficacy). As such, we collected one eDNA sample per pond and per time point, resulting in a total of 12 eDNA samples. To detect potential genetic contamination in the field, we collected four sample blanks by filtering mineral water on site: three associated with treatment evaluation (summer 2022, winter 2023, and summer 2023), and one associated with winter sampling in 2024.

During each sampling occasion, we conducted a composite eDNA sampling approach to account for the heterogeneous distribution of eDNA particles in lentic systems (Mayne et al., 2024), following Brys et al. (2021). Using a telescopic sampling pole from the pond shore, we collected 0.5 L of surface water, and transferred it into a decontaminated reservoir. We then repeatedly moved 5-10 m along the shoreline, collecting additional 0.5 L samples until the entire pond perimeter had been traversed. From this homogenised merged water sample, we pumped water through an enclosed disc filter equipped with an integrated 5-μm glass-fibre prefilter and a 0.8-μm polyethersulfone (PES) membrane (NatureMetrics, Surrey, UK) filter, using a sterile silicone tube attached to a peristaltic pump. We halted filtration upon clogging and recorded the total filtered volume. To prevent genetic contamination, we decontaminated all reusable equipment (i.e. sampling pole, reservoir, and peristaltic pump) after each sampling event using 2% Virkon S (Antec DuPont, Suffolk, UK). All filters (n = 16) were sealed and stored at −21°C until molecular analyses.

We conducted analytical processing of eDNA samples in the Research Institute for Nature and Forest (INBO) laboratories in Geraardsbergen, Belgium. We extracted DNA from filters using the Qiagen DNeasy extraction kit and purified it with the Qiagen DNeasy PowerClean Cleanup Kit. *Xenopus laevis* eDNA concentrations were quantified following the protocol described in Everts et al., (2025a). Briefly, we used the primer-probe protocol developed by Secondi et al. (2016), and adapted this for a droplet digital PCR (ddPCR) platform. Although these primers amplify all species within the *Xenopus* genus and are therefore not species-specific (Secondi et al., 2016), they reliably reflect *X. laevis* presence in Belgium, where it is the only *Xenopus* species known to occur (Pauwels et al., 2023; Everts et al., 2025a). Droplet digital PCR enables absolute quantification of target DNA concentrations through extensive sample partitioning and Poisson-based statistical inference (Huggett et al., 2016), and is comparatively robust to PCR inhibitors that are abundantly present in lentic environments resembling the sampled ponds (Harper et al., 2019). Previous studies have demonstrated that amphibian eDNA concentrations quantified *via* ddPCR in similar ecosystems correlate strongly with species densities (Everts et al., 2022). For each sample, we normalised eDNA concentrations by dividing by the total volume of water filtered (in L) to account for variation in filtrate volume among samples. We analysed each sample in triplicate, calculated *X. laevis* eDNA concentrations following Everts et al., (2022), and averaged replicate measurements per sample. To monitor potential contamination and analytical failure within the laboratory pipeline, we included five no-template controls and two positive controls containing 10 pg/µL of DNA extracted from *X. laevis* tissue.

#### Post-treatment visual evaluation & pH measurements

We checked pitfall traps at the inside of the fence for *X. laevis* presence on a daily basis for a period of two weeks following quicklime application. We conducted visual inspection by walking along the perimeter of the pond, and searching for *X. laevis* carcasses. To monitor recovery of the pH level of the water after treatment, we measured pH in all three treated ponds on January 24^th^ (days after finishing the treatment), February 2^nd^ (one week after finishing the treatment), and February 24^th^ (roughly a month following treatment), using a WTW MultiLine 3430 Portable Digital Multiparameter.

## Results

Fyke netting prior to quicklime application confirmed the low anticipated conservation value of the treated ponds. The vast majority of captured individuals were *X. laevis* individuals, while only small numbers of native fish (*Pungitus pungitius*), amphibians (*Lissotriton vulgaris*, *Mesotriton alpestris*, *Pelophylax* sp.), and invertebrates (including *Dytiscus marginalis*) were found. More specifically, we captured 8 *X. laevis* individuals in Pond 1 (7 adults, 1 subadult), 318 in Pond 2 (20 adults, 89 subadults, 50 larvae), and 398 in Pond 3 (80 adults, 63 subadults, 56 larvae). The measured pH remained high (> 9 Pond 1, > 11 for Pond 2, > 10 for Pond 3,) for almost a month after treatment (Table 1). By February 24^th^ 2023, the pH of all three ponds had decreased to pre-treatment conditions (i.e. below 8).

**Table 1.** Water pH measurements after quicklime treatment: days after finishing the treatment (January 24^th^), one week after treatment (February 7^th^), and roughly a month after treatment (February 24^th^).

|  | 24/01/2023 | 07/02/2023 | 24/02/2023 |
| --- | --- | --- | --- |
| Pond 1 | 11.91 | 11.16 | 7.96 |
| Pond 2 | 9.14 | 9.11 | 7.96 |
| Pond 3 | 11.93 | 10.88 | 7.95 |

No *X. laevis* eDNA was detected in either field or lab negative controls, and positive controls indicated no failures had occurred during the analytical workflow. Water volume filtered was on average 1.3 (± 0.70 standard deviation) litres. In summer 2022, prior to quicklime treatment, *X. laevis* eDNA concentrations were highest in Pond 2 (53.9 copies / L), followed by Pond 3 (28.1 copies / L) and Pond 1 (15.9 copies / L) (Fig. 4a). Eight weeks following quicklime treatment (winter 2023), all ponds lower mean *X. laevis* eDNA concentrations compared to the preceding summer (Fig. 4b), resulting in complete non-detection in Pond 2 (corresponding to a 100.0% reduction relative to the winter level 2024), a concentration of 0.1 copies / L in Pond 1 (80.1% reduction), and 5.8 copies / L in Pond 3 (a limited reduction of 3.5%). Similar reductions in *X. laevis* eDNA concentrations were also observed in summer 2023, with *X. laevis* eDNA concentrations of 5.8 copies / L in Pond 1 (63.9% reduction), 24.8 copies / L in Pond 2 (54% reduction relative to the summer level 2022), and 5.5 copies / L in Pond 3 (80.6% reduction) (Fig. 4a).

**Figure 4.**
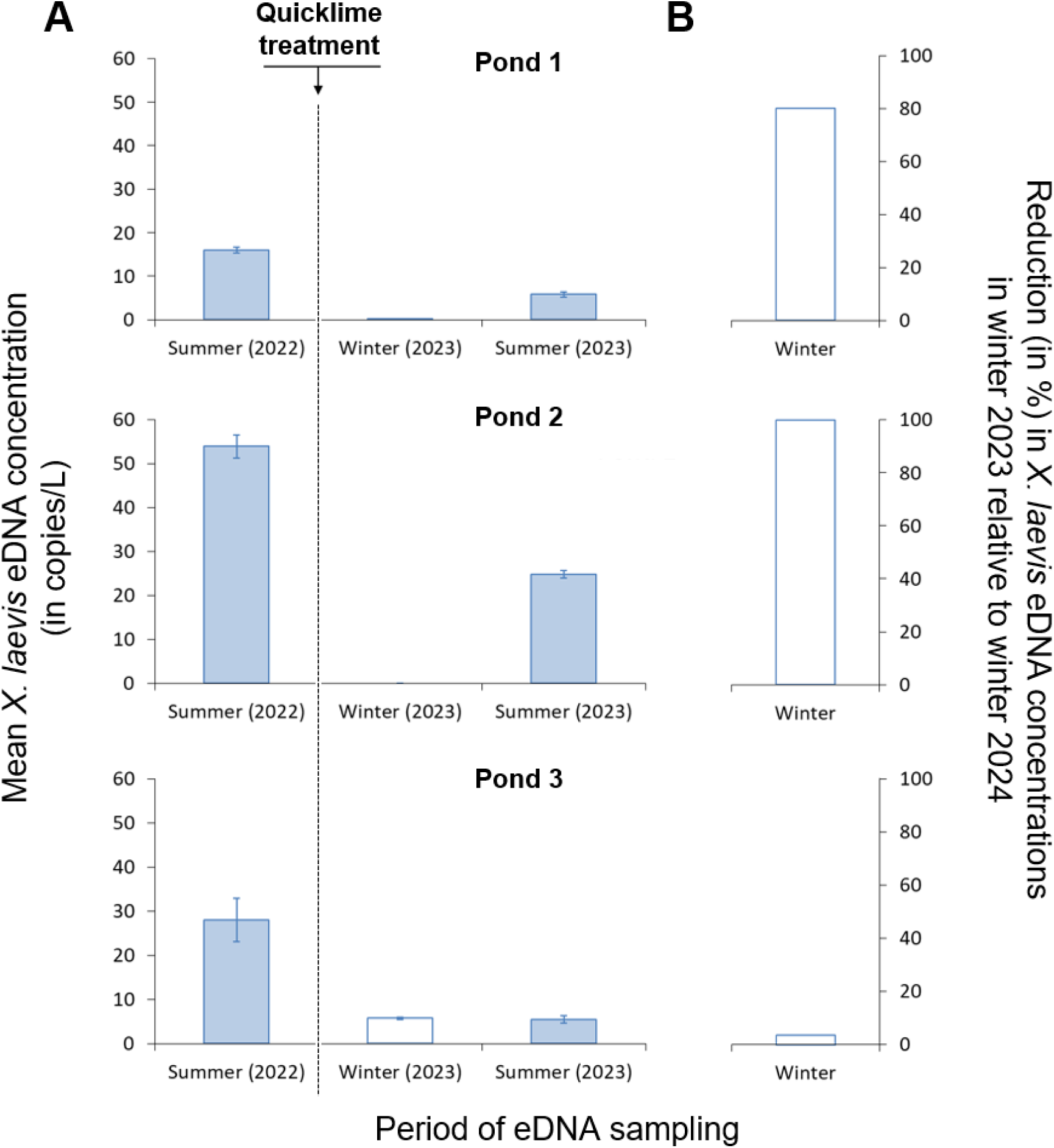
Evaluation of quicklime effectiveness in eradicating *X. laevis* in three ponds in Belgium. (**a**) Mean (and standard deviation) of *X. laevis* eDNA concentrations across three technical replicates measured before quicklime application (summer 2022), two weeks after quicklime application (winter 2023), and the summer following quicklime application (summer 2023). (**b**) Reduction (in percentage) of *X. laevis* eDNA concentrations in winter 2023 compared to the concentration in winter 2024 (a year following quicklime application) as a conservative estimate of quicklime effectiveness.

Visual post-treatment inspection demonstrated the effectiveness of the treatment. Hundreds of dead *X. laevis* post-metamorphs were observed near the shore of the three ponds. Specifically for Pond 3, 84 dead adult female individuals were counted. No *X. laevis* individuals were captured with the pitfall traps inside the fence.

## Discussion

### The efficacy of quicklime for eradicating X. laevis

Quicklime has been used as a high-intensity management tool in aquatic systems (Clearwater et al., 2008), including the removal of sea urchins in marine ecosystems (Christie et al., 2024) and non-native fishes from ponds and lakes (Britton et al., 2008; Abdel-Fattah, 2011; Van Kleef et al., 2015). Our results extend this evidence base by providing the first preliminary indication that quicklime can also be applied as an effective eradication treatment for non-native amphibians, by inducing mass mortality of *X. laevis* populations in small to medium-sized ponds (Witmer et al., 2015; Hegan, 2014; Tingley et al., 2017; Spikmans et al., 2022). As such, our work highlights the potential relevance of quicklime application for eradication or reproductive removal of pondscape invaders with aquatic life stages.

*Xenopus laevis* is particularly vulnerable to quicklime application as it is primarily aquatic in all life stages (Measey et al., 2012). Although decreasing water levels prior to quicklime application and subsequent extreme pH conditions may trigger burrowing behaviour, lime particles and alkaline porewater can penetrate superficial sediment layers, potentially causing lethal exposure even for individuals that temporarily seek refuge in the pond bottom (Collas, 2012; Boyd, 2017; Lazur et al., 2002). However, upwelling groundwater may locally dilute alkaline porewater and thereby reduce the lethality of quicklime in pond sediments. Quicklime efficacy can further be jeopardised when amphibians react to dropping water levels by leaving the water. During our intervention, such dispersal may have been limited by cold winter conditions (daily mean temperatures around 0 °C), whereas mass emigration in response to treatment is more likely during warmer periods. This underscores the importance of combining quicklime treatment with fencing and pitfall trapping, not only for amphibians but also for other mobile pondscape invaders such as crayfish and some invertebrates (Table 2). Although pitfall traps were installed to intercept dispersing individuals, not one *X. laevis* individual was captured on the pond-facing side of the fence. Rather than indicating treatment effectiveness, this absence can reflect limited *X. laevis* movement under the cold winter conditions during the intervention period.

**Table 2.**
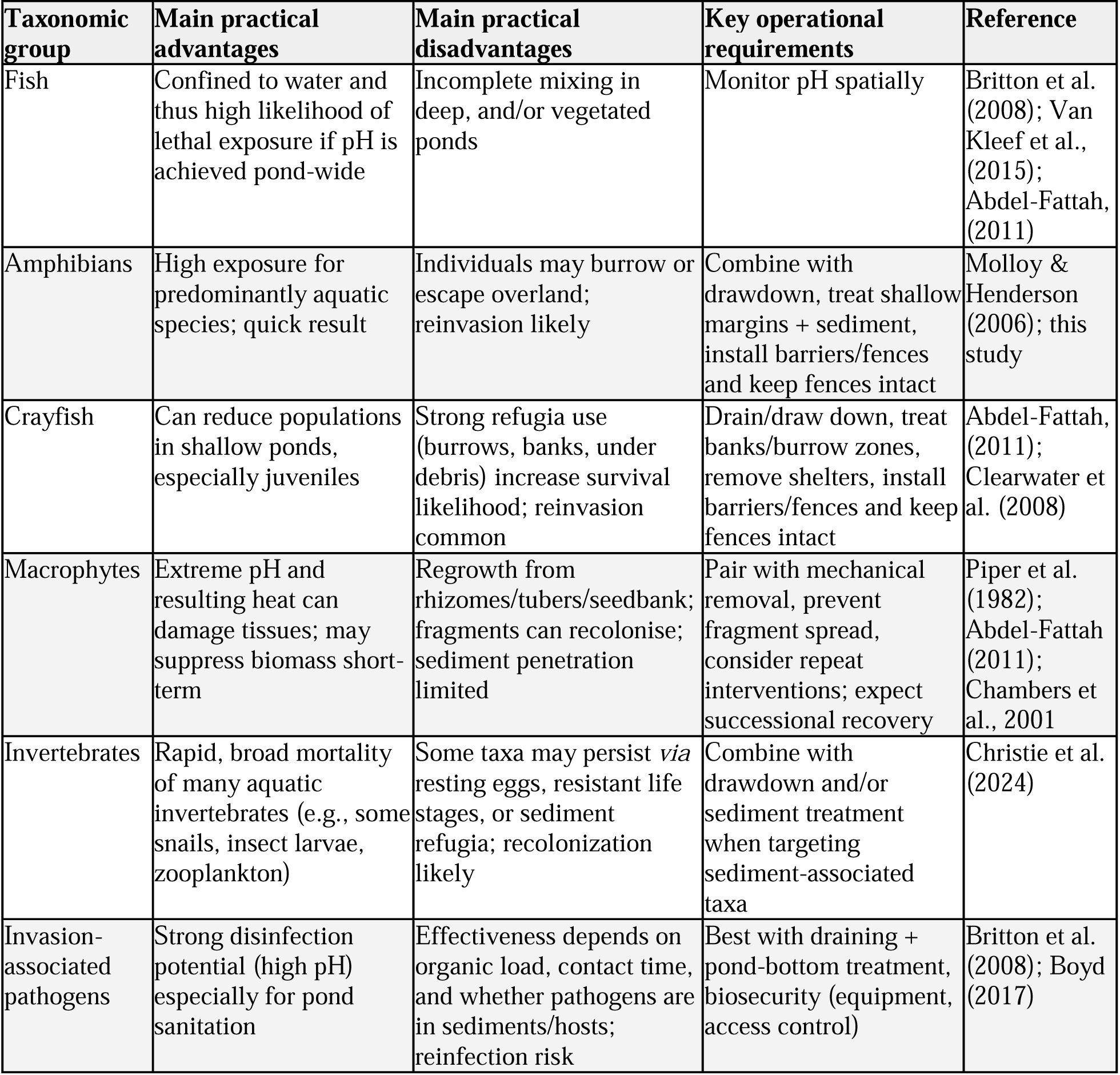
Overview of the main practical advantages, disadvantages, and operational requirements for some major taxonomic groups for which quicklime application can be a viable eradication measure.

| Taxonomic group | Main practical advantages | Main practical disadvantages | Key operational requirements | Reference |
| --- | --- | --- | --- | --- |
| Fish | Confined to water and thus high likelihood of lethal exposure if pH is achieved pond-wide | Incomplete mixing in deep, and/or vegetated ponds | Monitor pH spatially | Britton et al. (2008); Van Kleef et al., (2015); Abdel-Fattah, (2011) |
| Amphibians | High exposure for predominantly aquatic species; quick result | Individuals may burrow or escape overland; reinvasion likely | Combine with drawdown, treat shallow margins + sediment, install barriers/fences and keep fences intact | Molloy & Henderson (2006); this study |
| Crayfish | Can reduce populations in shallow ponds, especially juveniles | Strong refugia use (burrows, banks, under debris) increase survival likelihood; reinvasion common | Drain/draw down, treat banks/burrow zones, remove shelters, install barriers/fences and keep fences intact | Abdel-Fattah, (2011); Clearwater et al. (2008) |
| Macrophytes | Extreme pH and resulting heat can damage tissues; may suppress biomass short-term | Regrowth from rhizomes/tubers/seedbank; fragments can recolonise; sediment penetration limited | Pair with mechanical removal, prevent fragment spread, consider repeat interventions; expect successional recovery | Piper et al. (1982); Abdel-Fattah (2011); Chambers et al., 2001 |
| Invertebrates | Rapid, broad mortality of many aquatic invertebrates (e.g., some snails, insect larvae, zooplankton) | Some taxa may persist *via* resting eggs, resistant life stages, or sediment refugia; recolonization likely | Combine with drawdown and/or sediment treatment when targeting sediment-associated taxa | Christie et al. (2024) |
| Invasion-associated pathogens | Strong disinfection potential (high pH) especially for pond sanitation | Effectiveness depends on organic load, contact time, and whether pathogens are in sediments/hosts; reinfection risk | Best with draining + pond-bottom treatment, biosecurity (equipment, access control) | Britton et al. (2008); Boyd (2017) |

Management evaluation confirmed the short-term effectiveness of quickliming for killing off *X. laevis*. Large numbers of dead *X. laevis* post-metamorphic individuals were observed during post-treatment visual surveys. Short-term suppression was further supported by preliminary eDNA evidence: eight weeks post-treatment, *X. laevis* eDNA concentrations were undetectable in Pond 2 (100% reduction), and present in low concentrations in Pond 1 (80.1% reduction) and Pond 3 (3.5% reduction) relative to the same ponds sampled during the same period in the following year (Fig. 4). Importantly, residual *X. laevis* eDNA signals in Pond 1 and Pond 3 do not necessarily indicate incomplete eradication, as carcasses can continue to release detectable DNA for prolonged periods (Kamoroff & Goldberg, 2018; but see Curtis & Larson, 2020 for contradicting evidence). Indeed, at least 84 deceased *X. laevis* individuals were observed from the shore of Pond 3 following quicklime treatment. Additionally, cold winter temperatures may have reduced microbial activity, leading to prolonged persistence of DNA particles (Jo & Minamoto, 2021). In addition, the gradual decline in pH from highly alkaline conditions toward neutral levels may initially have limited DNA degradation, with degradation rates increasing through time once pH normalises again (Jo, 2026). Specifically for Pond 3, the detection of non-zero *X. laevis* eDNA concentrations, and thus limited apparent reduction, may be attributable to post-treatment pumping of water from the Douvebeek into the pond by farmers for irrigation purposes, which could carry *X. laevis* eDNA without indicating the presence of live individuals. These interacting effects, together with the lack of a more appropriate baseline eDNA signal (e.g. immediately before treatment during winter 2023) and insufficient knowledge on how *X. laevis* phenology influences eDNA concentrations at its northernmost invasion front (Everts et al., 2021), indicate that the obtained eDNA concentrations should be interpreted with caution as a potential guideline for determining eradication success in this specific case (but see Everts et al., 2022 for successful application elsewhere).

Despite evidence for near-complete eradication, *X. laevis* eDNA concentrations increased to considerable levels in all three ponds by the following summer, yet were substantially lower than before treatment (Fig. 4). Given that eDNA concentrations can roughly reflect species densities (Yates et al., 2019), particularly for amphibians in small, closed systems (Everts et al., 2022), these results indicate at least partial population recovery of *X. laevis* within the same year. Whether this recovery resulted from recolonisation from nearby waterbodies (Everts et al., 2025a), from survival of a small number of individuals, or both remains uncertain.

Importantly, quickliming may also suppress amphibian pathogens by reducing or eliminating their key reservoir host (Hossack et al., 2023) and by exposing the aquatic environment to elevated pH levels and extreme temperatures for prolonged periods (Chew et al., 2024). In the present case, chytrid fungi and ranavirus, which are commonly associated with *Xenopus* species, are therefore expected to be adversely affected by this treatment (Britton et al., 2008). In our study we did not assess pathogen dynamics, yet the effect of quicklime on pathogens merits further research.

Our findings suggest that quicklime can generate substantial mortality and rapid short-term reduction of *X. laevis* in invaded ponds, although the exact proportion of the population removed was not quantified. More generally, quickliming may be applicable to other non-native invasive amphibians, especially those with prolonged aquatic residence and pond-breeding habits, such as American bullfrogs (*Lithobates catesbeianus*), water frogs (*Pelophylax* spp.), as well as non-native species from other taxonomic groups (Table 2).

### Practical considerations

From an operational perspective, quickliming is best viewed as a high-intensity, whole-pond intervention, that requires careful preparation and planning to maximise effectiveness while limiting non-target impacts (Table 2). A typical liming operation ideally involves fencing and reducing the water volume to a manageable size, followed by intensive netting and trapping to remove as many remaining individuals as possible prior to treatment (Clearwater et al., 2008). Importantly, pond draining and fencing can be expensive and logistically and practically challenging, particularly in difficult-to-access habitats (e.g. pump installation and crane placement, reluctance of site owner permissions), groundwater-fed ponds (e.g. draining), or not clearly delineated water bodies (e.g. fence placement). While seasonal dry periods (e.g. summer droughts) can be leveraged to reduce draining effort and ease fencing, this timing may not align with the period during which the target species is most vulnerable to the quicklime application, such as during dormant stages. At the same time, applying quicklime during the species-specific period of greatest vulnerability may coincide with sensitive periods for native species (e.g. breeding), thereby creating a conservation dilemma. Because quicklime is non-selective, its use is generally unsuitable in high-quality habitats or water bodies supporting protected species, unless temporary removal prior to treatment and reintroduction or translocation are feasible options. It is most defensible in small, enclosed (often artificial) ponds where complete disruption of the aquatic community is considered acceptable (Clearwater et al., 2008). Following treatment, carcass accumulation can generate secondary water-quality effects (e.g., oxygen depletion, odour nuisance) and attract scavengers, making carcass removal and disposal a potentially important logistical consideration. Additionally, risk management during and immediately after treatment is essential, as extreme pH conditions can pose acute hazards to wildlife and potentially to domestic animals (Abdel-Fattah, 2011; Clearwater et al., 2008; Van Kleef et al., 2015). The use of quicklime also presents health and safety risks to operators (Table 2).

In addition to these constraints, the success and acceptability of quickliming depend strongly on site-specific hydrology and physicochemical conditions. Even small ponds may be connected *via* overflow pipes, ditches, or episodic flood events. Alkaline discharge to nearby waters can pose substantial environmental risks. Moreover, the quicklime dose required to achieve lethal pH is context-dependent, varying with water volume, organic matter load, sediment characteristics, groundwater inflow, and buffering capacity. Consequently, pH should be monitored at multiple locations, times, and depths to avoid untreated refugia and incomplete exposure.

Importantly, the effects of adding calcium-based alkaline materials to aquatic systems extend beyond short-term pH elevation. When quicklime is added to the water, subsequent reactions with dissolved carbon dioxide result in the formation of calcium carbonate (CaCO). This process alters water chemistry and sediment composition, with consequences that may persist long after the initial pH perturbation has diminished. Calcium carbonate influences hardness, buffering capacity, and ion balance in freshwater systems. These changes affect nutrient availability, microbial processes, and species composition, potentially reshaping ecosystem function (Clair and Hindar, 2005). Such effects are particularly pronounced in acidic, oligotrophic systems, where baseline buffering capacity is low and biological communities are adapted to nutrient-poor, low-calcium conditions. In these environments, even modest additions of calcium can shift relationships among primary producers, alter decomposition rates, and influence the structure of food webs. The persistence of CaCO in sediments may therefore represent a long-term effect, decoupled from the immediate objective of pH modification. (Bellemakers et al., 1994).

### Advantages and limitations

Deciding whether or not to use quicklime as a biological invasion eradication approach requires weighing the advantages and limitations in each specific context (Table 3). Quickliming, particularly in combination with fencing and pond drawdown, can be costly (Clearwater et al., 2008) and entails substantial collateral impacts, potentially resulting in the complete removal of all aquatic organisms present. Moreover, quickliming shares several limitations with conventional eradication approaches (e.g. incomplete removal), while introducing additional constraints (e.g. highly non-selective). For these reasons, it has been argued that conventional chemical methods may remain preferable to quickliming in certain management contexts (Van Kleef et al., 2015).

**Table 3.**
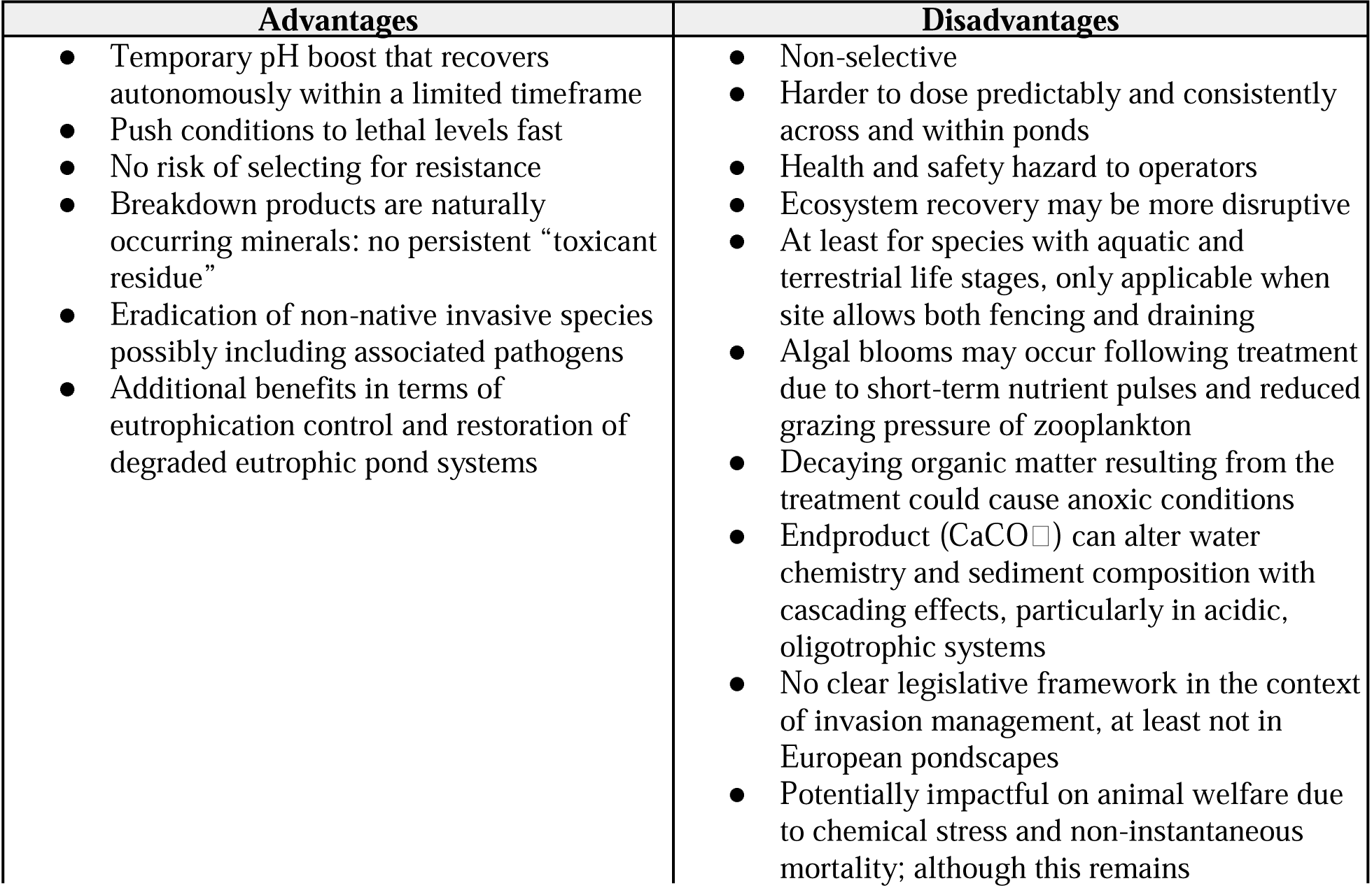
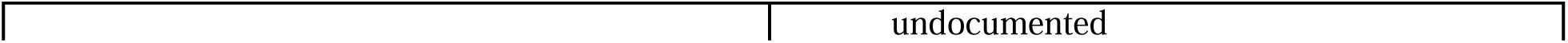
Advantages and disadvantages of quicklime application compared to conventional chemical eradication methods (e.g. rotenone, antimycin A, herbicides) for eradicating non-native invasive species in ponds.

However, quicklime also has advantages that cannot be overlooked (Table 3). Especially in the earliest invasion stages of highly invasive non-native species potentially carrying co-introduced pathogens in heavily degraded and eutrophic ponds, as illustrated with the French-Belgian *X. laevis* invasion here, quickliming may provide a viable alternative to eradication with other chemicals. In such contexts, it can be more effective than conventional trapping, whose effectiveness typically correlates with target species population sizes (Cooke & Beddington, 1984). Moreover, this non-selectiveness may also represent an advantage, as whole-system treatments have the potential to eliminate co-introduced pathogens, even when pathogens are not explicitly targeted (Brooks et al., 2020; Chew et al., 2024), as well as the other non-native invasive species tending to dominate pondscapes (Brönmark & Hansson, 2002) that are unintentionally removed. Furthermore, in biologically impoverished ponds, quickliming provides additional benefits in terms of eutrophication control by binding phosphates and nitrogen and can therefore be considered a method of ecological restoration (Prepas et al. 2001; Yang et al. 2025).

Quicklime application is not a silver bullet, but rather an approach to consider in the management toolbox of pondscape invaders, where a combination of methods (e.g. quicklime combined with fencing and trapping) is most desirable (Simberloff, 2021). Given the substantial collateral impacts associated with quickliming, we recommend its use primarily within early-detection rapid-response programmes in eutrophic ponds, where drastic interventions at a limited number of sites may achieve complete and permanent removal of the target species and possibly associated pathogens, rather than as a tool for long-term containment or mitigation. We recommend consideration of the whole-ecosystem context upon application, including both positive and negative, biotic and abiotic, non-target effects.

### Way forward

Knowledge on the use of quicklime for the eradication of aquatic invasive species and their associated pathogens remains limited (Clearwater et al., 2008). Here, we demonstrate the potential of quicklime application to combat invasive amphibians and other pondscape invaders, providing preliminary evidence of its applicability as a management tool. To achieve its full operational use, further research is needed into the cost-effectiveness compared to other management techniques, species-specific vulnerabilities and potential non-target impacts (Brooks et al., 2020), treatment timing (Vimercati et al., 2017; Lazur et al., 2002), resistance development (Clearwater et al., 2008), specific conditions (e.g. target species population size, species’ phenology, water temperature, habitat structure, CaO particle size) that favour or hamper the effectiveness of quicklime application (Brooks et al., 2020; Strand et al., 2020), effectiveness in removing associated pathogens (Britton et al., 2008), welfare impacts on target species, and target species behavioural responses (e.g. burrowing, leaving the waterbody) that may affect quicklime outcomes (Clearwater et al., 2008; Christie et al., 2024). Lab trials and small-scale field experiments are required to help reach these goals (Strand et al., 2020). Moreover, the relative effectiveness of quicklime compared with slaked lime (Ca(OH)) for eradicating pondscape invaders remains untested, particularly given that slaked lime induces pH elevation without the additional thermal shock associated with quicklime application. Although applying quicklime or slaked lime aimed at eliminating nuisance species is frequently practiced by fish farmers in parts of Europe (Wurts & Masser, 2013; Boyd, 2017; Lazur et al., 2002), a clear legislative and ethical framework governing its use in natural or semi-natural waterbodies is largely lacking. Consequently, methodological, regulatory, and ethical issues associated with drastic liming approaches such as quicklime application must be addressed before it can be used more widely to control other pondscape invaders.

## Acknowledgements

We thank Natuurwerk vzw for carrying out and reporting all management efforts, Sander Devisscher for data management, and Hanne Danneels and Nico De Regge for assisting in sample collection.

## Ethics statement

All field interventions were conducted with permission of the landowners and under the oversight of the competent authorities. Implementation followed safety and animal welfare guidance to minimise unnecessary suffering and non-target impacts. Investigations by the Research Institute for Nature and Forest (INBO) were performed under a generic exemption on the Flemish Species Decree, permit numbers ANB/BL-FF/V19-00209-212 and 21-202358.

## Funding

This study was partly supported by the European Union’s Horizon Europe HORIZON-CL6-2024-BIODIV-01 project “GuardIAS - Guarding European Waters from IAS”, under grant agreement no. 101181413.

## Notes

### Competing Interest Statement

The authors have declared no competing interest.

